# Dissociating stimulus encoding and task demands in ECoG responses from human visual cortex

**DOI:** 10.1101/2025.04.23.648016

**Authors:** Zeeshan Qadir, Harvey Huang, Müge Özker, Daniel Yoshor, Michael S. Beauchamp, Kendrick Kay, Dora Hermes

## Abstract

Brain responses to sensory stimuli depend on the specific task performed by the observer. Previously, fMRI measurements in ventral temporal cortex (VTC) showed that BOLD responses to visual input scale with task demands. This scaling of BOLD responses is thought to be driven by the engagement of cognitive functions required for the task. To better understand how these cognitive task demands influence neural activity in VTC, we measured electrocorticography (ECoG) responses to images of varying contrast in two human participants during either a fixation or image categorization task. ECoG high frequency broadband activity (>70 Hz) showed that local neuronal responses increased with image contrast as expected for sensory encoding, and were scaled by task demands approximately 200 ms after stimulus onset. In contrast, ECoG low frequency activity in the alpha/beta range (8-28 Hz) was insensitive to image contrast and showed larger decreases with increasing task demands. These results indicate that high frequencies in VTC reflect both the encoding of sensory inputs and task demands, similar to prior BOLD response measurements, whereas low frequency oscillations represent task demands, not sensory inputs per se. In line with the interpretation that low frequency oscillations reflect pulsed inhibition, we speculate that a power decrease in low frequency oscillations amplifies neural activity in VTC in support of visual task demands.

## 1 Introduction

Tasks can be viewed as sets of objectives that influence how the brain processes sensory visual input. Objectives vary across tasks and may engage different cognitive functions, including spatial attention, perceptual decision making, reading, and memory (for detailed review on tasks, see Kay et al. 2023). Through these objectives, tasks modulate sensory neural responses to meet required cognitive demands. Some of these task effects, like spatial attention, are well established: studies have investigated how spatial attention influences neural spiking activity in the visual cortex. Early models of spatial attention first captured how attention may gate visual input by suppressing irrelevant information (Moran and Desimone 1985) and then evolved to additionally capture attentional enhancement of relevant information through various gain models (McAdams and Maunsell 1999a; Reynolds et al. 2000; Reynolds and Chelazzi 2004; Reynolds and Heeger 2009). Spatial attention, however, is only one of the many cognitive functions that visual tasks may involve (Kay et al. 2023). Tasks through these cognitive functions thus shape visual processing, and comprehensive theories of vision must account for how tasks modulate neural activity in the visual cortex.

One theoretical framework proposed to capture task modulations that involve cognitive processes beyond spatial attention is flexible top-down modulation (Zhang and Kay 2020). The flexible top-down modulation framework posits that experimental tasks scale responses in relevant regions in order to process specific stimuli and meet task demands. Task demands are defined here as the cognitive processes required for achieving given task objectives. This framework was shown to be effective in explaining task modulation of fMRI BOLD responses for some cognitive processes (Kay et al. 2015; Kay and Yeatman 2017). As there are many different cognitive processes (Corbetta and Shulman 2002; Gold and Shadlen 2007; Heekeren et al. 2008; Nobre and Gresch 2025), it is important to understand to what extent the neurophysiological mechanisms that support top-down modulation of visual areas follow common principles.

Top-down signaling in visual areas is captured in different signal features in field potential recordings, including high and low frequency components. Broadband signals (spanning across many frequencies and often extracted in a high frequency broadband (HFB) range of 70-170 Hz) are thought to capture local neuronal processing and often correlate with population firing rate (Miller et al. 2009a; Manning et al. 2009; Ray and Maunsell 2011; Hermes et al. 2017). An interesting question is what the temporal dynamics of top-down signaling are. Tasks might induce sustained effects on visual responses that last the entire duration of stimulus presentation, suggesting sustained maintenance of top-down signaling (Silver et al. 2007; schematically shown in Figure 1*b-ii*). Or, tasks might induce short transient effects suggestive of brief control signals that shift cognitive states (Yantis et al. 2002; schematically shown in Figure 1*b-iii*). Temporally resolved HFB signals can capture the temporal dynamics of these task modulations thus revealing how these visual responses are modulated during tasks.

**Figure 1:**
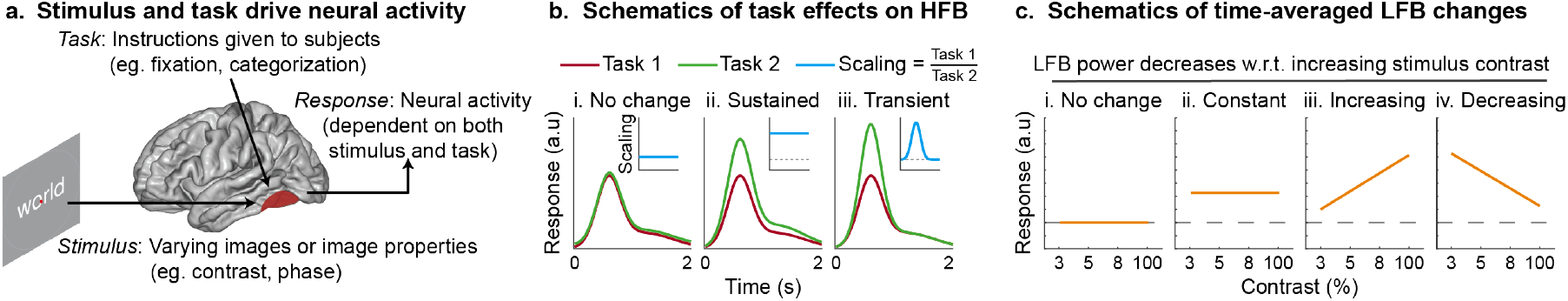
Schematics of stimulus and task driven visual responses. (a) Schematic showing that recorded neural activity is dependent on both stimulus and task. (b) Schematics of (i) a lack of task effects with no differences in the HFB responses across two tasks, (ii) sustained task effects with constant scaling, and (iii) transient scaling effects. Cyan traces in the insets indicate scaling factor (ratio of responses of Task 2 over Task 1). The dashed line indicates no scaling. (c) Schematics showing possible changes for time-averaged LFB power decrease with varying stimulus contrast. Scenarios include (i) no change, (ii) constant change due to general task engagement independent of stimulus, (iii) largest change for high contrast stimulus, or (iv) largest change for low contrast stimulus which may induce the highest demands during an image categorization task (Kay and Yeatman 2017). The dashed line indicates no change with respect to the prestimulus baseline.

Besides HFB, top-down influences on local processing have also been captured in low frequency (LFB) oscillations in the alpha range (8-12 Hz; Jensen and Mazaheri 2010; Schalk 2015) as well as in the alpha-beta range (8-24 Hz; Bauer et al. 2014; Bastos et al. 2015; Richter et al. 2017). Large alpha oscillations during rest are proposed to inhibit cortical firing, and decreases in the power of these oscillations may therefore reduce inhibition. Power changes in these lower frequencies have been related to distractor inhibition, gain modulation, as well as gating during spatial attention (Zhigalov and Jensen 2020; Bonnefond and Jensen 2024). Recently, an EEG-fMRI study (Griffiths et al. 2019) suggested that alpha/beta power decreases reflect general increases in stimulus information processing during three different tasks involving visual perception, auditory perception, and visual memory retrieval, indicating a broad modulatory role of these oscillations across cognitive tasks. We therefore investigate whether alpha/beta power changes relate to the effects of task demands previously captured by the flexible top-down modulation framework. If alpha/beta power changes indeed reflect task demands, we expect a larger decrease for more difficult low contrast conditions (Figure 1*c-iv*).

To better understand how tasks modulate neural responses in visual areas, we recorded electrocorticography (ECoG) in two human participants as they engaged in two different tasks with varying cognitive task demands. We took advantage of a paradigm from a previous fMRI study where subjects perform fixation and categorization tasks while they look at words and faces of varying contrast (Kay and Yeatman 2017). This previous study proposed the flexible top-down modulation framework to account for observed task effects on stimulus driven fMRI BOLD responses. By re-using this paradigm, we were able to verify that BOLD changes are related to neural processing observed in ECoG HFB responses, extract temporal profiles of top-down modulation in ECoG HFB responses, and test the hypothesis that alpha/beta power plays a general role across experimental tasks in tracking task demands. Our analyses confirm that post-stimulus HFB power is jointly modulated by the stimulus and task demands, tracking previously reported BOLD changes, and also show additional transient temporal dynamics related to task modulation. On the other hand, post-stimulus LFB power decreases across the alpha/beta range uniquely reflected task demands and were not driven by the stimulus. These results support the idea that alpha/beta power changes may capture task modulation and play a general role in scaling local neural activity to meet task demands across experimental tasks.

## 2 Methods

### 2.1 Subjects

The experimental protocol was approved by the Institutional Review Board of the Baylor College of Medicine. Two patients (F/30 (Sub-1) and F/52 (Sub-2)) with medically refractory epilepsy, who had intracranial electrode arrays (AdTech) implanted for clinical epilepsy monitoring to localize seizure foci before surgical treatment, consented to participate in the study. A third patient also consented, but had a lesion near the visual word form area (vWFA) and had difficulty seeing the stimuli and so was excluded from the study.

Electrode locations were estimated based on the operative sketch, and are displayed on the MNI (Montreal Neurological Institute)-152 template brain to provide a schematic overview (Figure 3*a**&b*). These are not meant to perfectly capture the anatomy, and we note that none of the analyses performed in this paper depend on detailed anatomical labeling. We grouped electrodes into those reflecting early visual areas (EVC, posterior to the posterior transverse collateral sulcus (ptCOS)) and those reflecting ventral temporal cortex (VTC, anterior to ptCOS).

### 2.2 Stimuli and Tasks

Subjects were presented with face and word stimuli, varying in contrast and phase coherence, with a central fixation dot that changed color every 600 ms. The stimuli and tasks were the same as used in the previous fMRI study (Kay and Yeatman 2017; Figure 2). The face stimuli (faces101:7) were made freely available without copyright restrictions from the lab of Dr. Kalanit Grill-Spector. Face stimuli were presented at contrast levels of 4%, 6%, 10%, and 100% whereas word stimuli were presented at contrast levels of 3%, 5%, 8%, and 100%. Both sets of stimuli were also presented at 0%, 25%, 50%, 75%, and 100% phase coherence levels. Other control stimuli consisting of houses, checkerboards, and polygons were also presented but were not considered in our analyses.

**Figure 2:**
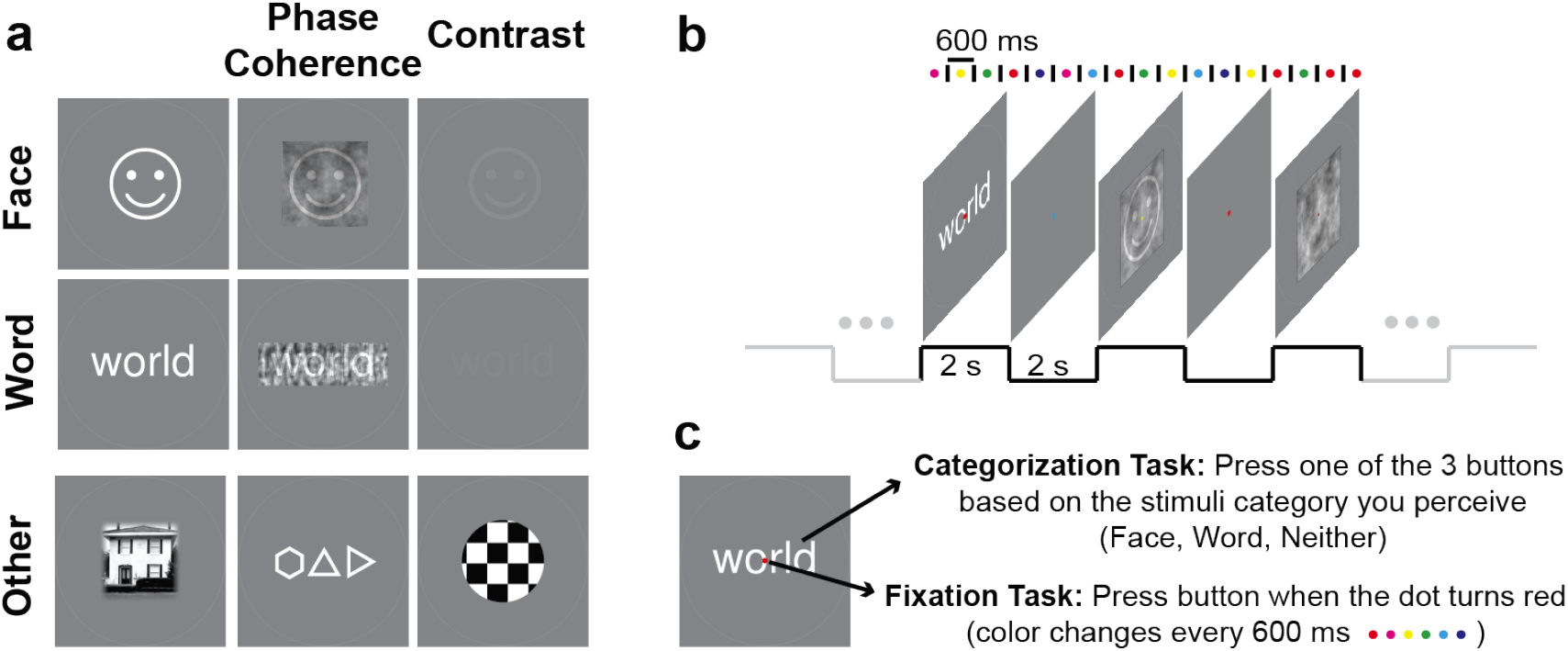
Experiment task design. (a) *Stimuli*. Stimuli included faces and words presented at different contrasts and phase-coherence levels, as well as full-contrast houses, polygons, and checkerboards. Faces shown here is for illustration purpose only. During the experiment pictures of human faces were shown. (b) *Run*. Each run consisted of all the stimuli interleaved with a blank screen, with each stimulus presented thrice in a random order. Notice that the fixation dot changed color independent of the change of stimulus. (c) *Tasks*. On a given run, subjects performed either a fixation or a categorization task.

**Figure 3:**
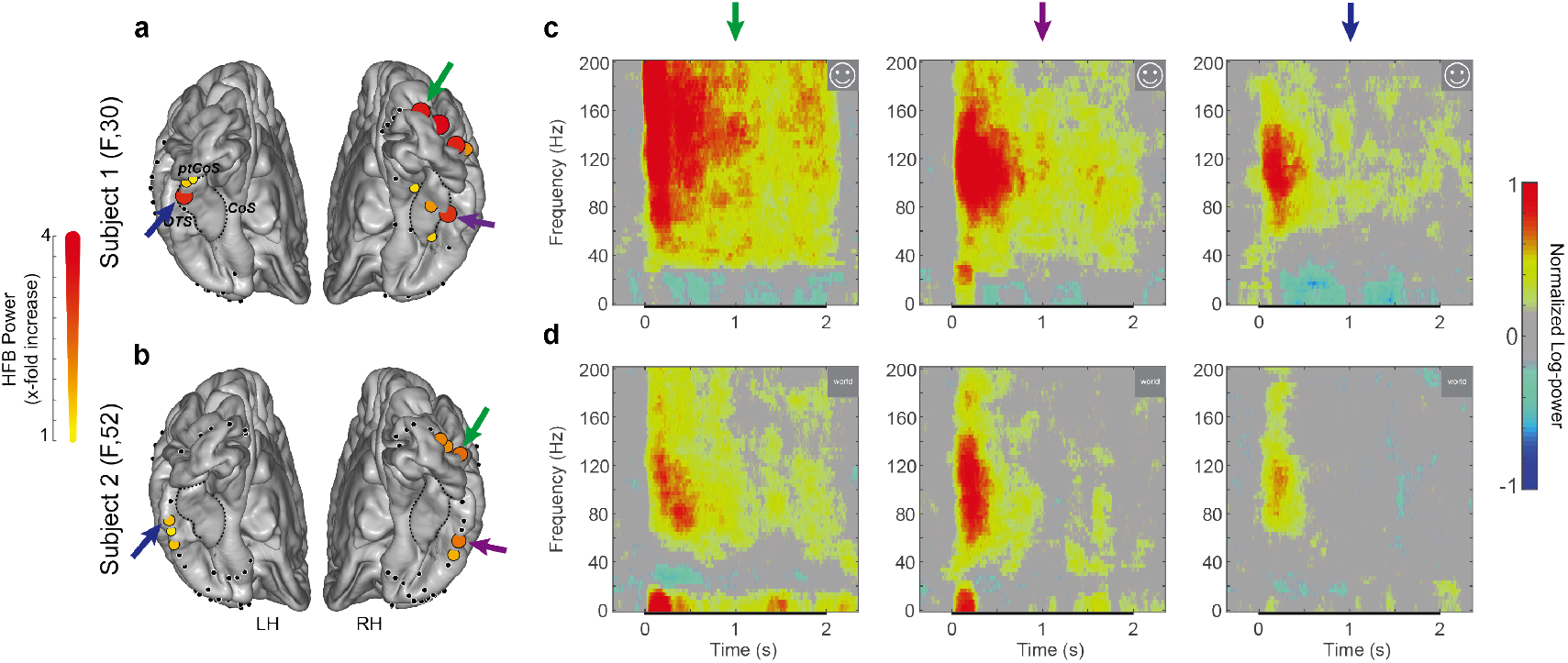
Electrode locations and spectral power changes. (a,b) ECoG electrode locations in Subject 1 (top) and Subject 2 (bottom). Visual-responsive electrodes are shown in colored circles; color and size of the circle indicates the intensity of time-averaged HFB power increase (0-400 ms, face & word responses at 100% contrast). (c,d) Time-frequency power (spectograms) for an EVC electrode (green arrow) and a RH VTC electrode (purple arrow), and a LH VTC electrode (blue arrow) in each subject. The inset shows the stimulus (face/word at 100% contrast) for which the plots were computed. Black bars on the time axis indicate duration during which stimulus was presented (0-2 s). *COS* : collateral sulcus, *ptCOS* : posterior transverse COS, *OTS* : occipital-temporal sulcus

In separate runs, subjects were asked to perform a fixation task and a categorization task. The fixation task is intended to minimize the influence of task demands on how the stimuli are processed. The categorization task is intended to probe how task demands influence sensory processing.

#### Fixation task

Subjects were asked to press a button whenever the fixation dot turned red. The dot changed color every 600 ms in a random order to one of six colors: magenta, red, yellow, green, cyan, and blue. Dot color changes were continuous (i.e., no rest) and the timing of the dot changes (every 600 ms) was independent of the presentations of the stimuli (every 2 s). This was intentional to minimize any potential interactions between the performance of the fixation task and responses to the stimuli. Red dot appeared for a total of 154 times in two runs (80 and 74) for subject 1 and 140 times (70 and 70) for subject 2.

#### Categorization task

Subjects were asked to press one of three buttons – word, face, or neither – based on the stimulus category they perceived. Subjects were instructed to respond within 4 s of stimulus onset (this coincided with 2 s of stimulus presentation plus 2 s of rest/blank presentation). The task was adjusted for the ECoG setting, and therefore differed slightly from the mini-block design used previously during fMRI (Kay and Yeatman 2017). During ECoG, each run consisted of 68 trials: 1 blank trial, 66 stimulus trials, and 1 blank trial. During each stimulus trial, the stimulus was presented for 2 s followed by rest (blank) for 2 s. Both subjects performed 2 runs of each task in the order of FCFC (F: fixation, C: categorization). Each stimulus (e.g., face at 3% contrast) was presented three times during each run giving a total of 6 trials for each combination of stimulus and task for each subject. The viewing distance for the experiment was approximately 60 cm, and the stimuli were approximately 2° tall and wide. A faint circle (10% opacity, 4.2° diameter) surrounding each stimulus was shown for subject 1 to help them realize when the stimuli were present, and this circle was presented identically for all stimuli. Both the tasks were roughly matched for the number of button presses: 140-154 times during fixation, 132 times during the categorization task. Unlike the previous fMRI study, we did not conduct a one-back task in the ECoG study.

### 2.3 ECoG recording

ECoG data were recorded at 2000 Hz from a Blackrock neural signal processor. Non-recording electrodes (as well as electrodes near the seizure onset zone) were identified based on clinical notes and were excluded from analysis. Raw signal from the remaining electrodes was first filtered (5th order Butterworth IIR notch) for line noise removal at 60 Hz and its harmonics. Common average re-referencing (CAR) was then performed to remove other electrical noise common to all electrodes. Data were reviewed and epochs with artifacts (sharp transients and interictal discharges) were removed.

### 2.4 ECoG time varying HFB and LFB responses

The signal from each electrode was bandpass filtered (2nd order Butterworth IIR), in frequency bins of 5 Hz; 70-170 Hz for high frequency broadband (HFB) and 8-28 Hz for lower frequency (LFB; alpha/beta) oscillations. These frequency ranges are consistent with previous studies reporting power fluctuations in HFB (Miller et al. 2009a; Manning et al. 2009; Ray and Maunsell 2011; Hermes et al. 2015) and LFB (Bauer et al. 2014; Griffiths et al. 2019; Yuasa et al. 2025). Then, the absolute value of the Hilbert transform was calculated and the geometric mean was taken across bands (to ensure that lower and higher frequency bands contributed relatively equally to the average). Next, the power of the signal was calculated and the signal was downsampled to 200 Hz. We calculated the x-fold signal change with respect to the prestimulus baseline power (e.g., power of 2 during baseline and power of 8 during stimulus is a 3-fold increase: 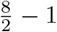). The prestimulus baseline was defined as 500 ms to 100 ms before stimulus onset, and baseline power was averaged across time and conditions using the geometric mean. The HFB or LFB signal changes are plotted as power increases or decreases with respect to 0 (no change).

Visually responsive electrodes were identified as electrodes with significant HFB power increases (*p <* 0.01 by right-tailed sign test, Bonferroni corrected for multiple comparisons across all electrodes) in the 0-400 ms time window post stimulus onset, pooling across all full contrast conditions. These visually responsive electrodes were classified based on anatomical location as reflecting early visual cortex (EVC; *n*_*Sub*−1_ = 4, *n*_*Sub*−2_ = 3) and ventral temporal cortex (VTC; *n*_*Sub*−1_ = 7, *n*_*Sub*−2_ = 5). Electrodes that were visually responsive, but not within these regions were not considered for our analysis (*n*_*Sub*−1_ = 8, *n*_*Sub*−2_ = 3). Similar as in other studies (Jacques et al. 2016), VTC electrodes were further classified into fusiform face area (FFA) and visual word form area (vWFA) electrodes based on their *d*^′^, calculated as

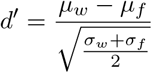

where *µ*_*x*_ is the average HFB response in 0-400 ms post stimulus onset, *σ*_*x*_ is the variance across 100% contrast trials, and *x* ∈ {*w* = *word, f* = *face*}. If *d*^′^ >= 1, then electrodes were marked as vWFA (*n*_*Sub*−1_ = 2, *n*_*Sub*−2_ = 3), whereas if *d*^′^ <= −1, then electrodes were marked as FFA (*n*_*Sub*−1_ = 4, *n*_*Sub*−2_ = 0). Other electrodes (−1 *< d*^′^ < 1) were considered to be non-selective VTC (*n*_*Sub*−1_ = 1, *n*_*Sub*−2_ = 2) electrodes.

### 2.5 ECoG HFB scaling factor

To quantify the effect of cognitive task demands, a scaling factor was computed. First, HFB responses (averaged across trials) were smoothed using a 200 ms moving average window. Then, the scaling factor was computed as the ratio of HFB response during the categorization task over HFB response during the fixation task for each condition. HFB responses that fell below baseline were set to 0.01 to avoid division by zero or negative numbers. We used cluster-based permutation testing that corrects for multiple comparisons (Maris and Oostenveld 2007) to determine time points that were significantly (*p* < 0.05 different across the two tasks. Non-significant scaling factors were set to 1 (indicating no scaling).

### 2.6 Average HFB and LFB power changes

In order to compare the power changes observed in this study with the BOLD percent signal changes reported in (Kay and Yeatman 2017), we time-averaged ECoG responses in a 1 second window post stimulus onset. HFB responses were averaged across a 0-1 s window to capture the bulk of the responses. LFB responses were averaged across a 0.25-1.25 s window that incorporated a slight temporal offset in order to minimize the effect of evoked responses.

### 2.7 Spectral visualization

To visualize spectrograms, we used a multi-taper time-frequency spectrum analysis (Bokil et al. 2010; 300 ms moving average window, 10 ms step size). Spectrograms for each condition were created by computing the median spectral response across trials. Baseline was calculated as the median across time (baseline window: 500 ms to 100 ms before stimulus onset) and conditions. Finally, spectrograms were log-normalized by computing the log and subtracting the baseline.

### 2.8 Principal spectral component analysis

Several studies have emphasized the importance of separating low frequency oscillations from the background 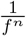 power (Miller et al. 2009b, 2014; Hermes et al. 2015, 2019; Donoghue et al. 2022; Yuasa et al. 2025; van Engen et al. 2026). When filtering for low frequency bands, the overall power may be affected by a combination of spectral broadband change plus additional oscillatory signals. As a control analysis, we computed the power spectral components (PSCs; Miller et al. 2009a) using principal component analysis (PCA). This data driven approach extracts components that explain the most variance across the power spectrum. In ECoG data, the first component often reflects broadband power and the second component often captures changes in lower frequencies (Miller et al. 2009a). The PCA ensures that these features are orthogonal and changes in one are uncorrelated with changes in the other. First, the power spectral density was computed in four bins of 500 ms corresponding to data from 500 ms prestimulus to 1500 ms post stimulus onset. Next, the power spectrum was flattened (across tasks, stimuli, and trials) and was then log-normalized by taking log of the power spectrum and subtracting the average power spectrum. PCA on the power spectrum produced PSCs in decreasing order of variance explained. The first two PSCs were then considered for further analysis to ensure that low and high frequency band changes capture different processes, rather than different spectral features of the same process.

### 2.9 Time-to-onset, time-to-peak, and duration

To calculate temporal response features, responses were pooled across face and word conditions for EVC electrodes. For selective VTC electrodes, responses to faces were pooled for FFA electrodes and responses to words were pooled for vWFA electrodes. Next, we used cluster-based permutation testing that corrects for multiple comparisons (Maris and Oostenveld 2007) to find time windows (within 0-1 s of ECoG response post stimulus) for which responses were significantly (*p* < 0.05) different compared to the prestimulus baseline for each task. *Time-to-onset* was then calculated as the initial time point of the first significant cluster and *duration* was calculated as the total length of all the significant time windows (i.e., significant clusters). Lastly, *time-to-peak* was calculated as the time at which the response was maximum for each condition, again, within 0-1 s of ECoG response post stimulus. Statistical analysis was performed by fitting a linear mixed-effects model that predicted time-to-onset, time-to-peak, and duration, independently, with fixed effects for stimulus contrast X task X ROI (EVC/VTC), and an intercept-only random effect for subjects.

### 2.10 Behavioral response

Reaction time and accuracy were computed during the experiment. For both tasks reaction time was calculated as the time taken by the subjects for their first button press from stimulus onset. For the fixation task reaction, the button presses within 1.2 s of dot change was considered a valid response as there was at least 1.2 s gap between two consecutive red dots. In case of multiple responses during this window, only the first response was considered. Subject 1 missed a button press 6 of the 154 times the dot changed to red, while subject 2 had 9 misses of 140 times. For the categorization task, in case of multiple responses only the first response was considered. If no response was recorded within 4 s after stimulus onset, the trial was considered as invalid and excluded from the behavioral analysis. Similar to the fixation task, in case of multiple responses only the first response was considered. Only one condition, face contrast at 4%, had two invalid trials in subject 1. All other conditions had at least one response from both subjects during categorization. Average *reaction times* were then calculated as the median of time to first button presses across all valid trials. Next, *response accuracy* during the categorization task was calculated as the percentage of correct responses across all valid trials. For the 0% and 25% phase coherence conditions, we considered the ‘neither’ response as correct.

## 3 Results

In order to understand how task demands affect the neurophysiological processing of visual inputs, two patients with ECoG electrodes implanted for epilepsy monitoring performed, in separate runs, a fixation task and a categorization task while they looked at words, faces and other stimuli (Figure 2). These stimuli were varied in contrast and in phase coherence. During the fixation task, subjects pressed a button when a central dot turned red, irrespective of the stimulus. As noted in the fMRI study (Kay and Yeatman 2017), the aim of the fixation task is to serve as a control task to minimize the influence of task demands on how sensory information is processed. We acknowledge that the fixation task does not necessarily prevent a general awareness of the stimulus. During the categorization task, subjects pressed a button to indicate whether the stimulus was a word, face, or neither, while still maintaining fixation on the central dot. Categorization required subjects to make a perceptual decision, which both subjects were able to do well (with 83-100% accuracy, Figure S1*b*). Both subjects were faster at categorizing high contrast stimuli compared to lower contrast stimuli (average reaction times for highest and lowest contrast stimuli in Sub-1: 0.85 s ± 0.05 SEM and 1.00 s ± 0.07 and in Sub-2: 1.65 s ± 0.10 and 1.84 s ± 0.07). Consistent with many psychophysiological experiments (Pins and Bonnet 1996; Barbur et al. 1998; Plainis and Murray 2000), the longer reaction times during categorization of lower contrast stimuli suggests that task demands are higher for low contrast compared to high contrast stimulus conditions. While subjects were overall faster to categorize face stimuli with higher phase coherence compared to faces with lower phase coherence, the differences in reaction times between phase coherence variations for word stimuli were smaller (Figure S1*b*). To best capture how neurophysiological processes change with task demands, we therefore focus the ECoG results on the responses to contrast variations and show results for phase coherence in the supplement (Figures S4, S5, S6*b*, S8*b*).

### 3.1 Visual stimuli and task demands induce independent HFB and LFB changes

Robust visual responses were measured in 19 electrodes, bilaterally implanted, across the two subjects (*n*_*Sub*−1_ = 11, *n*_*Sub*−2_ = 8). These electrodes were grouped into early visual cortex (EVC) and ventral temporal cortex (VTC) electrodes, located anterior and posterior to the posterior transverse collateral sulcus (ptCOS), respectively (Figure 3*a**&b*). The visual stimuli induced HFB power increases as well as LFB power decreases. Spectrograms, in example EVC and VTC electrodes in each subject, show that HFB increases were highest during the first 0-1 s of stimulus presentation, and gradually reduced for the remainder of the stimulus duration of 2 s (Figure 3*c**&d*). LFB power decreases were relatively smaller and most evident after ∼0.2 s, after evoked response related power changes are resolved. These HFB and LFB changes are consistent with previous ECoG studies of visual cortex (Hermes et al. 2015; Jacques et al. 2016).

Recent studies emphasized how analyses of oscillatory power changes can sometimes be confounded by broadband shifts in power (also referred to as aperiodic shifts; Miller et al. 2009b, 2014; Hermes et al. 2015, 2019; Donoghue et al. 2022; Yuasa et al. 2025; van Engen et al. 2026). To understand the effects of top-down modulation on visual processing, it is therefore necessary to carefully dissociate broadband and low frequency oscillations. In order to test whether the HFB and LFB power changes capture different aspects of the same underlying neurophysiological signal or capture distinct sources of neurophysiological changes, we performed a spectral principal component analysis (PCA). The spectral PCA naively extracts spectral features without preselection of a specific frequency range, and will result in separate components if stimuli and task conditions have different effects on low and high frequencies (for details, see Miller et al. 2009a). The spectral PCA identified a broadband component (1-200 Hz) as the first principal spectral component (PSC) and a low frequency component as the second PSC (Figure S2*c**&d*). The low frequency component ranges from ∼8-28 Hz and captures alpha and beta oscillations, indicating that in our experiment these low frequencies (8-28 Hz) fluctuate together. These two PSCs in broadband and low frequency range are consistent with previous studies (Miller et al. 2009a; Sabra et al. 2020) and reveal that these two components likely represent different neurophysiological processes. These observations from our data-driven PCA analysis justify our choices of how to quantify HFB and LFB power changes: we compute HFB increases by filtering ECoG responses in 70-170 Hz and LFB decreases by filtering responses across the alpha/beta range in 8-28 Hz (Figures 4, 5, 6). Supplementary analyses of power changes based on the spectral PCA components and separate alpha and beta bands show similar results (Figures S6*c**&d*, S8).

**Figure 4:**
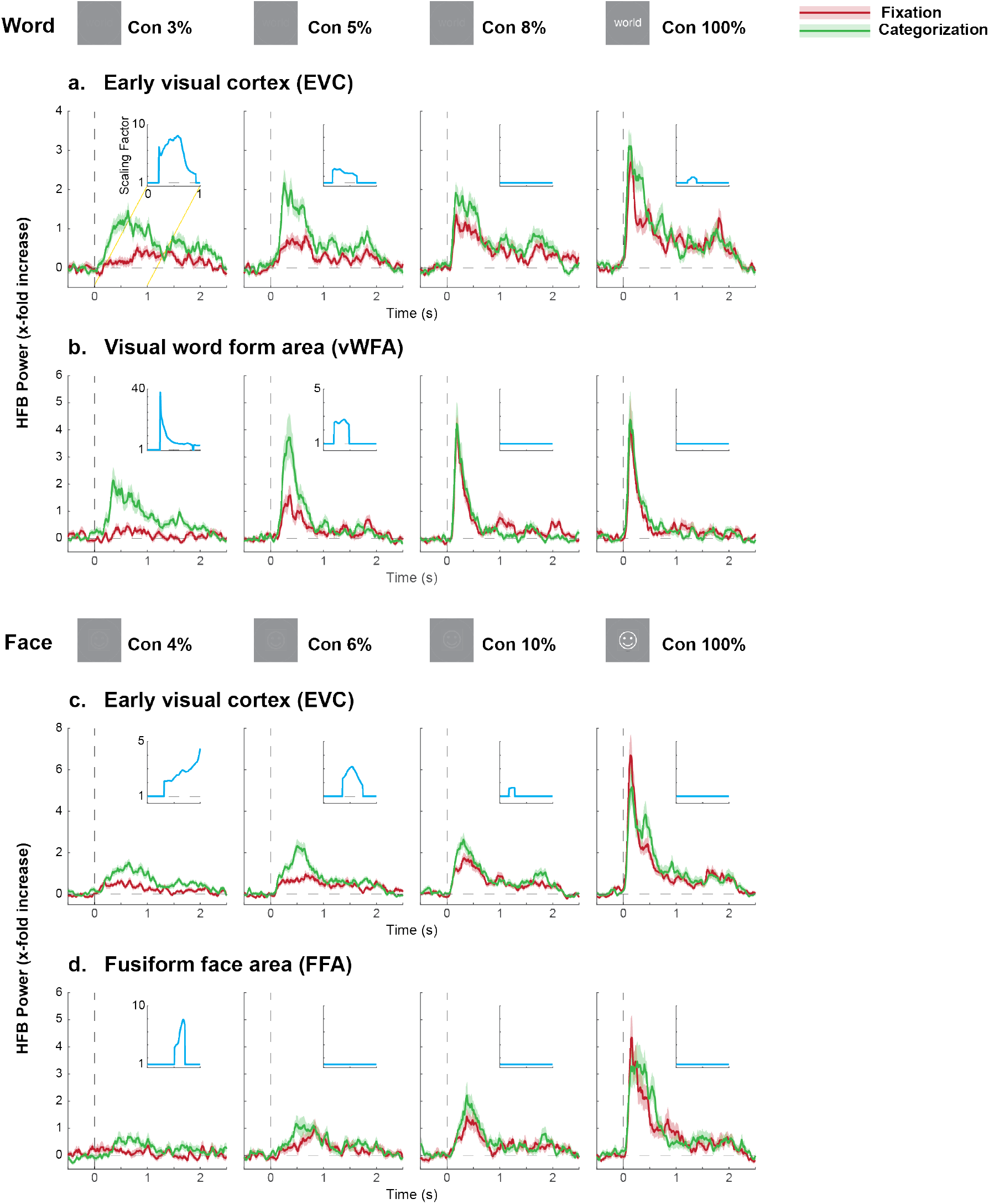
Stimulus and task effects in ECoG HFB. ECoG HFB power traces during fixation (red) and categorization (green) with increasing stimulus contrast (word/face, shown on the top). The average HFB power (solid) was calculated by geometric averaging over electrodes and trials. The shaded region corresponds to the 68% confidence interval. The horizontal dashed lines indicate baseline and the vertical dashed lines indicate the stimulus onset. The inset in each plot corresponds to the scaling factor calculated as the ratio of HFB power of the categorization over the fixation task. Cyan traces deviated from 1 (grey dashed line) when scaling effects were significant (p<0.05; cluster-based permutation test). Dashed lines at 1 indicate no scaling.

**Figure 5:**
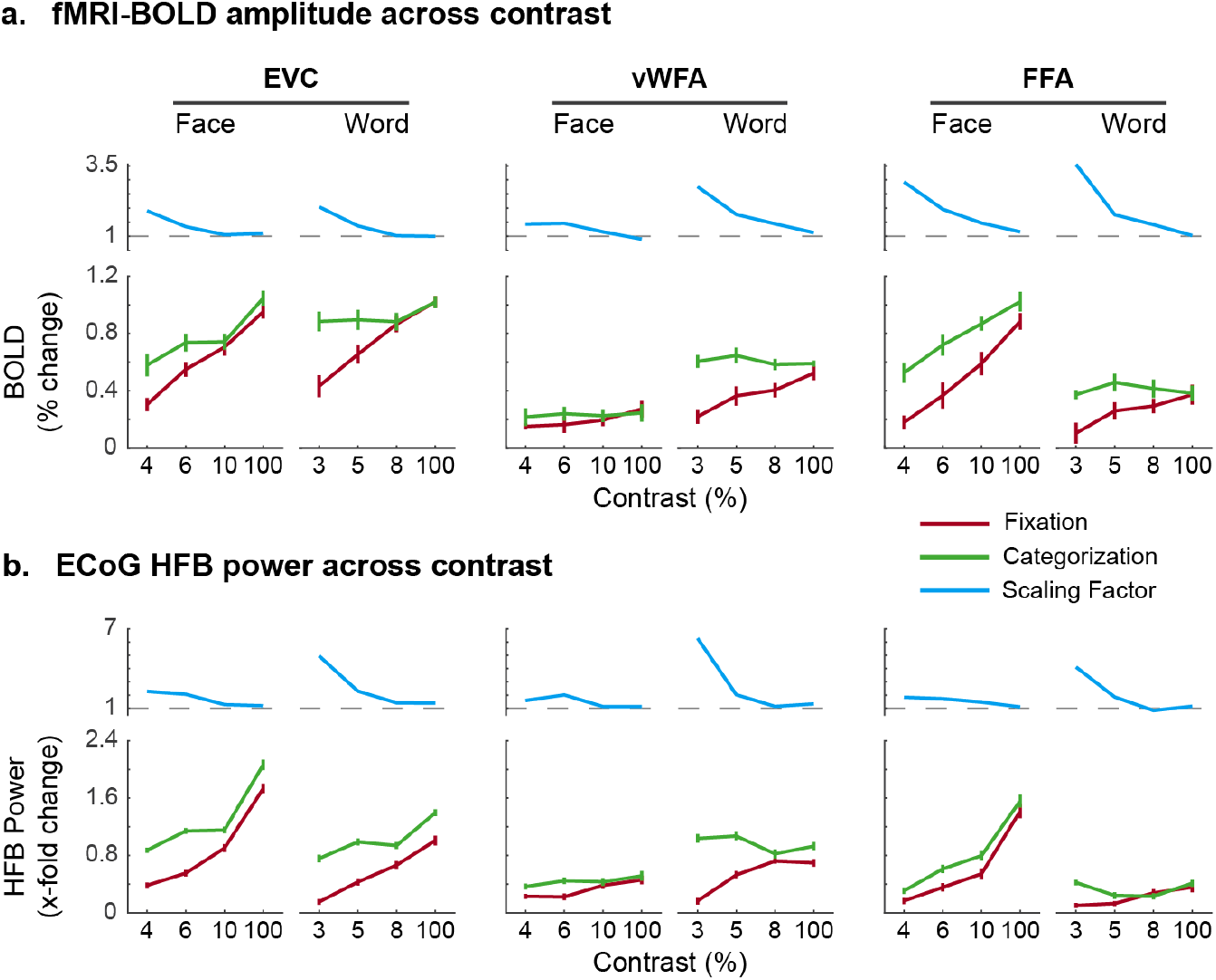
Time-averaged ECoG HFB responses match fMRI BOLD amplitudes. (a) fMRI BOLD responses reproduced from (Kay and Yeatman 2017) for varying stimulus contrast for EVC (left) vWFA (middle) and FFA (right). (b) ECoG HFB responses time-averaged in 0-1 s time window. Cyan traces show scaling factors with horizontal dashed lines at 1 indicating no scaling. Error bars indicate 68% confidence interval.

**Figure 6:**
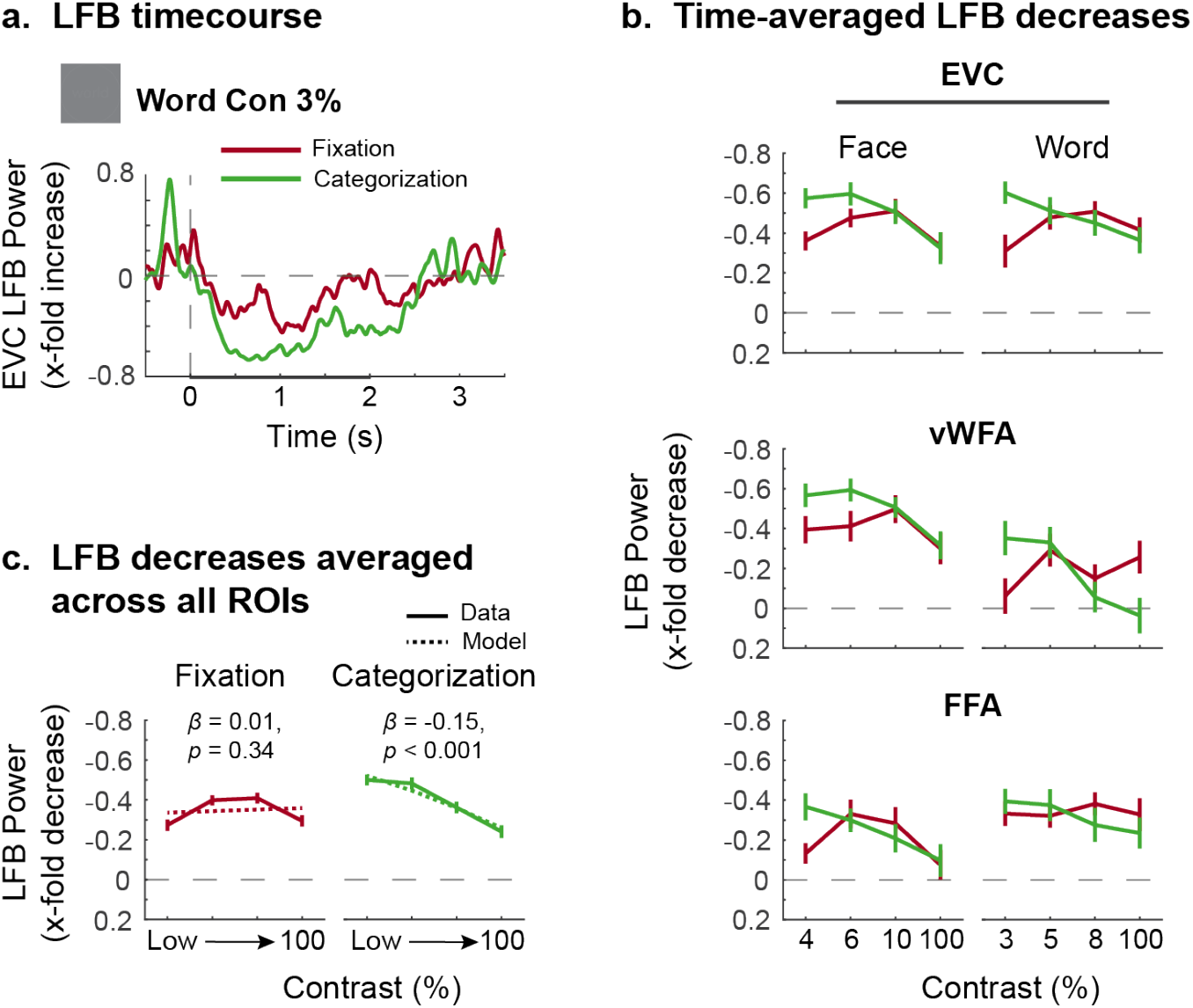
Time-averaged ECoG LFB responses reflect task demands. (a) LFB time course shown for 3% word contrast averaged across all EVC electrodes, as an example. Black bar on the time axis indicates duration during which stimulus was presented (0-2 s). (b) LFB power decreases time-averaged in 0.25-1.25 s time window. (c) LFB decreases for faces and words combined and averaged across all the EVC, vWFA, and FFA electrodes. Dotted line indicates the best linear fit with slope *β*. Error bars indicate 68% confidence interval. Note: the vertical axis in b & c is inverted to emphasize LFB decreases.

### 3.2 Task demands transiently increase HFB responses in a stimulus specific manner

HFB responses can reveal the timing of the effects of task demands on neural activity. As described in the previous fMRI study (Kay and Yeatman 2017), BOLD responses were shown to scale with task demands. To test whether this scaling affects the entire response duration or has a transient effect (represented in schematics in Figure 1*b*), we calculated time-varying HFB responses. We show that the HFB responses encoded stimulus features and were affected by the task as follows.

First, HFB responses encoded stimulus features in a way that replicates previous ECoG studies (e.g. Henrie and Shapley 2005; Dubey and Ray 2020; Groen et al. 2022). Increasing stimulus contrast resulted in larger HFB responses throughout EVC, FFA and vWFA, regardless of the task (Figure 4). Moreover, the time-to-onset and the time-to-peak both decreased with increasing stimulus contrast (time-to-onset: *β* = −114.16, *S*.*E*. = 17.34, *p <* 10^−5^, time-to-peak: *β* = −120.58, *S*.*E*. = 27.65, *p* < 10^−5^; Figure S3). The temporal dynamics of the HFB response further show that high contrast words and faces induced large initial responses for 1 s following the stimulus onset, which was then followed by a smaller sustained response until 2 s, after which the HFB power returned to the baseline. These delays and onset transients are consistent with previous modeling and empirical studies (Zhou et al. 2018, 2019).

Second, the HFB responses were affected by the task performed by the subject. During categorization, HFB responses were typically larger compared to fixation. For example, responses to word stimuli at 5% contrast during fixation peaked at less than 1-fold HFB increase compared to baseline, whereas during categorization, responses increased by more than 2-fold (Figure 4*a*, second panel). These increases were particularly pronounced for lower contrast stimuli, and may be indicative of increased task demands required to categorize these lower contrast stimuli (Kay and Yeatman 2017). Note, these response patterns might be interpreted using a contrast-gain model (Reynolds et al. 2000, but see Discussion 4.3).

To quantify the temporal profile of this task-related scaling, we calculated time-resolved scaling factors as the ratio of the HFB response during categorization over fixation (Figure 4 insets, cyan traces indicate significant scaling (*p* < 0.05, cluster-based permutation test)). Task-related scaling effects were transient; the scaling factor had an onset around 0.2 s (range: 0.190-0.600 s), mean: 0.360 s ± 0.040 SEM) and typically lasted for <1 s (with one exception in Figure 4*c*, left panel). These scaling effects were observed for faces and words in the EVC (Figure 4*a**&c*). Word selective electrodes in the VTC (visual word form area, vWFA) showed strong, transient scaling for low contrast word stimuli (Figure 4*b*). Similarly, face selective electrodes in the VTC (fusiform face area, FFA) showed strong, transient scaling for low contrast face stimuli (Figure 4*d*). Scaling was also present in EVC electrodes for face and word stimuli during phase coherence, although these effects were not observed in VTC (Figure S4, S5). The latter might be due to the fact that we did not expect categorization of stimuli for lower (0% and 25%) phase coherence conditions (see Methods 2.10), whereas we did expect categorization of stimuli at lowest contrast conditions which typically showed the highest scaling effects.

While the previous fMRI study showed that BOLD responses were increased during categorization, our results further reveal that increases in neural activity are not present during the entire response period, but are visible most strongly during ∼0.2-1 s. Thus, ECoG provides us with temporal information of these task-related increases, i.e., task demands are more prominent during the transient period with a delayed onset and are not sustained for the entire response period.

### 3.3 ECoG HFB responses match fMRI BOLD

Previous studies that compared BOLD and field potential measurements reported a strong correlation between the fMRI BOLD response and the ECoG HFB changes (Hermes et al. 2015, 2017; Jacques et al. 2016). Our results are consistent with these findings. The previous fMRI study (Kay and Yeatman 2017) showed that BOLD responses were increased during categorization compared to fixation, and this increase was the strongest during low contrast conditions with high task demands (Figure 5*a*). We compared these BOLD signal changes from the previous study to the time-averaged HFB responses from 0-1 s (Figure 5*b*). Time-averaged scaling factors were calculated as the ratio of the time-averaged HFB responses during categorization over fixation. The modulatory scaling effects are remarkably similar across the two measurement modalities, with lower contrast conditions showing large increases in task-related scaling. Therefore, both modalities indicate that increased response scaling is observed with higher cognitive task demands.

### 3.4 Low frequency alpha/beta power decreases encode task demands

To investigate how alpha/beta power changes relate to changing task demands previously captured by the flexible modulation framework, we extracted LFB power in 8-28 Hz. LFB power decreased after the initial evoked response during both tasks, with larger decreases observed during categorization (Figure 6*a*). Moreover, the magnitude of the time-averaged LFB power changes was different for the two tasks. During the fixation task, LFB power decreased in a comparable magnitude from baseline across low and high contrast conditions (Figure 6*b*, red lines). However, during the categorization task, LFB power decreases varied across stimulus contrast, with ∼3-fold larger decreases during the lowest contrast stimulus compared to the highest contrast stimulus (Figure 6*b*, green lines). The stronger LFB decreases when categorizing low contrast face and word stimuli was consistently observed across EVC, FFA and vWFA.

In order to formally test how the two different tasks affected LFB power, we averaged LFB power across stimulus types and fit a first order polynomial relating contrast to the response (Figure 6*c* dotted lines). On the one hand, the line fit for the fixation task does not have a slope that is significantly different from 0 (*β* = 0.01, *p* = 0.34; similar to schematics in Figure 1*c-ii*), which confirms that the LFB decreases during the fixation task do not encode the stimulus variations. Instead, the constant decrease in LFB power may suggest an automatic reduction in LFB induced by a general engagement of the visual system. On the other hand, the line fit for the categorization task shows a significant negative slope (*β* = −0.15, *p* < 0.01; similar to schematics in Figure 1*c-iv*), indicating that LFB decreases during categorization are reduced with increasing stimulus contrast. Whereas task demands are relatively constant during the fixation task (where the subject continuously monitors the color of the fixation dot), the categorization task is most difficult in low contrast conditions and easiest during high contrast conditions. During the categorization task, LFB decreases, therefore, appears to reflect task demands. Overall, this pattern of LFB decreases suggests that visual input induces a constant reduction in LFB and additional variations in the amount of LFB decreases reflect task demands.

## 4 Discussion

Task demands influence how visual inputs are processed by the brain. We measured ECoG signals in human visual areas to characterize neural processing of visual inputs during two different tasks (a fixation task and a categorization task). We found two distinct neurophysiological signals - broadband high frequency power and low frequency alpha/beta oscillations - that encoded visual inputs and task demands. As expected, HFB responses encoded stimulus contrast and increased with higher contrast. HFB responses additionally exhibited modulation by task demands in terms of task-related scaling effects, which were most pronounced during difficult categorization conditions. Extending beyond what was previously shown in fMRI BOLD, we found that task-related scaling in local neuronal activity is transient, starting ∼0.2 s post stimulus onset and typically lasting < 1 s. On the other hand, LFB responses were not driven by stimulus contrast and therefore do not appear to encode stimulus properties per se. Rather, LFB responses reflected task demands, showing larger decreases for more difficult categorization conditions, selectively following task modulations consistent with the flexible top-down modulation framework.

### 4.1 Low frequency power reflects task demands

In the fixation task, LFB power decreases were relatively comparable across different contrast conditions. Conversely, in the categorization task, LFB power decreases were stronger for low contrast conditions. How do we reconcile these different results? LFB power has previously been interpreted as being sensitive to some aspects of visual input: some studies found that alpha power could be used to decode visual input (Chen et al. 2023), while others did not (Griffiths et al. 2019). One study detailed how alpha power decreases were spatially tuned to visual inputs with relatively large receptive fields while the subjects performed a similar fixation task (Yuasa et al. 2025). Alpha power in a neuronal population may therefore be sensitive to input to the population receptive field. However, we suggest that the level of alpha decreases may reflect task demands rather than the visual input to the receptive field per se. This interpretation is consistent with laminar recordings in macaque V1 that showed that alpha power decreased when an unattended visual stimulus was presented compared to when an auditory stimulus was presented, but decreases were much larger when subjects paid attention to the visual stimulus (Bollimunta et al. 2011). These results are also consistent with previously reported LFB decreases related to attentional demands (Osaka 1984; Ray and Cole 1985; Jensen and Mazaheri 2010; Babu Henry Samuel et al. 2018). Task demands therefore seem to regulate the magnitude of the LFB decrease.

Large low frequency alpha and beta oscillations at rest have been proposed to reflect pulsed inhibition of local neuronal firing (Jensen and Mazaheri 2010; Miller et al. 2012; Schalk 2015). According to this framework, pulsed inhibitory inputs suppress the membrane potential to inhibit local neuronal firing. Decreases in LFB power reduce inhibition in task-relevant regions, resulting in attentional enhancement of responses to attended objects (Zumer et al. 2014). The larger decreases in LFB power during difficult stimulus categorization conditions may reflect a reduction of inhibition that facilitates local neuronal computation — for example, upregulation of sensory-related signals in VTC may help inform decision-related processes performed in the intraparietal sulcus (Kay and Yeatman 2017). EEG studies have indeed linked low frequency power decreases to a state of optimal excitability in the underlying neuronal population (Rajagovindan and Ding 2011). High alpha power, conversely, may inhibit neuronal computation and serve as a gate for feedforward flow (Zhigalov and Jensen 2020). Overall, we suggest that decreases in alpha and beta power may facilitate the contribution of local neuronal processing to the performance of different visual tasks.

### 4.2 High frequency broadband response scaling is transient

The ECoG recordings enabled us to estimate the time window during which task demands scale neural responses. The HFB response increases during difficult categorization conditions (compared to fixation) were observed transiently from ∼0.2-1 s post stimulus onset. The scaling effects thus do not start with response onset nor do they last for the duration of the stimulus presentation. These results are also consistent with previously reported attentional modulation of HFB responses (Davidesco et al. 2013; Martin et al. 2019). These transient scaling effects are in line with the brief control signals postulated to shift cognitive states rather than a continuous signal actively maintaining the modulated state (Yantis et al. 2002). Indeed, the hypothesis of brief control signals is consistent with our prior observation (Kay and Yeatman 2017) that the amount of scaling is specific to the stimulus, in a way that appears to be related to the task being performed by the subject (e.g. evidence accumulation for perceptual decision making).

The fact that the scaling is observed relatively late supports the idea that the scaling is the result of feedback inputs to VTC. This feedback can arise from several areas. Feedback inputs to visual areas during the same categorization task were previously modeled as inputs from the intraparietal sulcus (Kay and Yeatman 2017). This previous study measured fMRI BOLD changes in parietal areas and found that parietal responses predicted the gain measured in the VTC. This is also supported by anatomical tracing and human diffusion MRI studies that show that parietal cortex and visual ventral temporal areas are directly connected (Felleman and Van Essen 1991; Distler et al. 1993; Webster et al. 1994; Markov et al. 2014; Yeatman et al. 2014; Takemura et al. 2016; Bullock et al. 2019; Panesar et al. 2019; Vinci-Booher et al. 2022). Further, TMS studies have also shown that parietal stimulation can change the excitability in early visual areas V1/V2 (Silvanto et al. 2009; Capotosto et al. 2009, 2012). While inputs from parietal cortex are one candidate for potentially driving the measured task modulations of visual cortex, frontal eye fields (Gilbert and Li 2013), and subcortical structures like the pulvinar (Desimone et al. 1995; Saalmann et al. 2012) and superior colliculus (Sommer and Wurtz 2004) likely also play a role in task modulation. We acknowledge the limited electrode coverage from 2 subjects in the current study, and that future electrophysiological measurements of such regions would be important to provide information on the neurophysiological basis of dorsal and ventral interactions during different visual tasks.

### 4.3 Task-related scaling is consistent with the flexible top-down modulation framework

The measured HFB responses increased with stimulus contrast and this contrast response function was modulated during the categorization task. Specifically, categorization increased HFB responses for low contrast conditions. These contrast response functions can shift in different ways with cognitive processing, and previous studies have proposed four different models that capture different types of shifts observed across various visual paradigms (Zhang and Kay 2020). First, an additive-shift model predicts an increase in neural activity of comparable magnitude across all contrast conditions (Luck et al. 1997; Kastner et al. 1999; Ress et al. 2000; Giesbrecht et al. 2006; Silver et al. 2007; Buracas and Boynton 2007; Murray 2008). Second, a response-gain model multiplies the responses by a gain factor, which results in increased gain for high amplitude responses to high contrast stimuli (McAdams and Maunsell 1999b), opposite of what we observe here. Third, a contrast-gain model predicts increased sensitivity by horizontally shifting the contrast response function and results in the largest response increases for mid-contrast stimuli (Reynolds et al. 2000). Lastly, a flexible top-down modulation framework predicts that the experimental task scales responses in relevant regions in order to process specific stimuli and meet certain task demands (Zhang and Kay 2020).

In both a contrast-gain and flexible top-down modulation framework, the response increases are particularly pronounced for lower contrast stimuli. However, as previously observed with fMRI BOLD responses (Kay and Yeatman 2017), the ECoG HFB responses to lower contrast stimuli sometimes exceed those of higher contrast stimuli. For example, HFB response for the 5% word contrast during categorization in EVC is higher than that for the 8% contrast during fixation (Figures 4*a*, 5*b*). Therefore, in a manner similar to the BOLD responses, the task driven ECoG HFB increases are not fully explained by a contrast-gain model that would predict a mere leftward shift in the contrast response function (Reynolds et al. 2000). The increased reaction times during low contrast conditions are consistent with the interpretation that task demands were highest when categorizing low contrast stimuli, and therefore the ECoG HFB scaling is consistent with the flexible top-down modulation framework (Kay and Yeatman 2017; Zhang and Kay 2020). Moreover, we observed that the power of low frequency oscillations changed according to the flexible top-down modulation, suggesting a potential mechanistic role of low frequency oscillations to support the observed HFB scaling effects.

### 4.4 Significance of task sampling for interpreting neural responses

Our findings demonstrate the importance of considering the task when evaluating and interpreting the relationship between visual inputs and neural responses. If we had sampled neural responses only while subjects performed the fixation task, we would have observed a constant LFB decrease irrespective of the stimulus presented. This may have led to an incorrect interpretation that LFB decreases were irrelevant to visual processing. On the other hand, if we had sampled neural responses only while subjects performed the categorization task, we might have come to the incorrect interpretation that LFB encodes stimulus contrast. Future research will need to sample a variety of different tasks in order to better characterize how task demands shape processing in the visual pathways.

## 5 Acknowledgments

The authors thank the patients and staff at the Baylor College of Medicine. The authors also thank Lupita Yanez and Kai Miller for their constructive feedback. Research reported in this publication was supported by the National Eye Institute of the National Institutes of Health under award numbers R01EY035533 (KK and DH), R01NS065395 (MSB) and R01EY023336 (DY). The content is solely the responsibility of the authors and does not necessarily represent the official views of the National Institutes of Health.

## 6 Additional information

### 6.0.1 Author contributions

Zeeshan Qadir: Software, Formal Analysis, Validation, Visualization, Writing-original draft, Writing-review & editing;

Harvey Huang: Software, Writing-review & editing;

Müge Özker: Investigation, Data Curation;

Daniel Yoshor: Investigation, Resources, Funding Acquisition, Writing-review & editing;

Michael S. Beauchamp: Methodology, Investigation, Resources, Funding Acquisition, Writing-review & editing;

Kendrick Kay: Conceptualization, Methodology, Software, Data Curation, Supervision, Funding Acquisition, Writing-original draft, Writing-review & editing;

Dora Hermes: Software, Formal Analysis, Supervision, Funding Acquisition, Writing-original draft, Writing-review & editing

## 7 Additional files

### 7.0.1 Data availability

To foster reproducible research, we will make the data and code publicly available upon final version via a permanent archive on the Open Science Framework and the code to reproduce the figures on Github.

## Supplementary Materials

**Figure S1:**
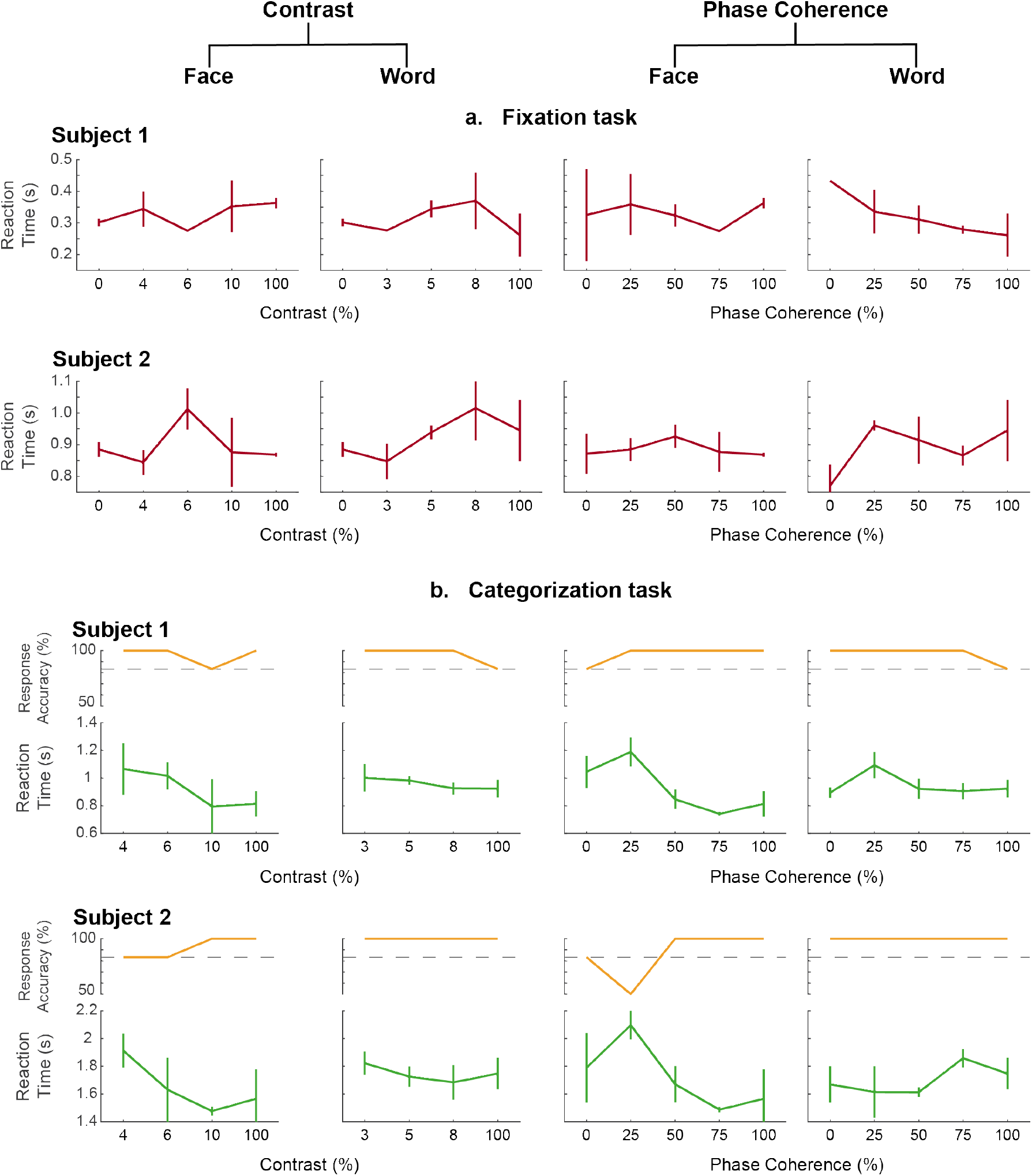
Behavioral response. The average reaction time for button press responses (median across trials) is shown for both the subjects during (a) fixation and (b) categorization task. Error bars indicate 68% confidence interval. Additionally, the response accuracy during the categorization task (b; orange traces) is shown for the two subjects with dashed lines indicating 83.33% accuracy (5 of 6 correct responses).

**Figure S2:**
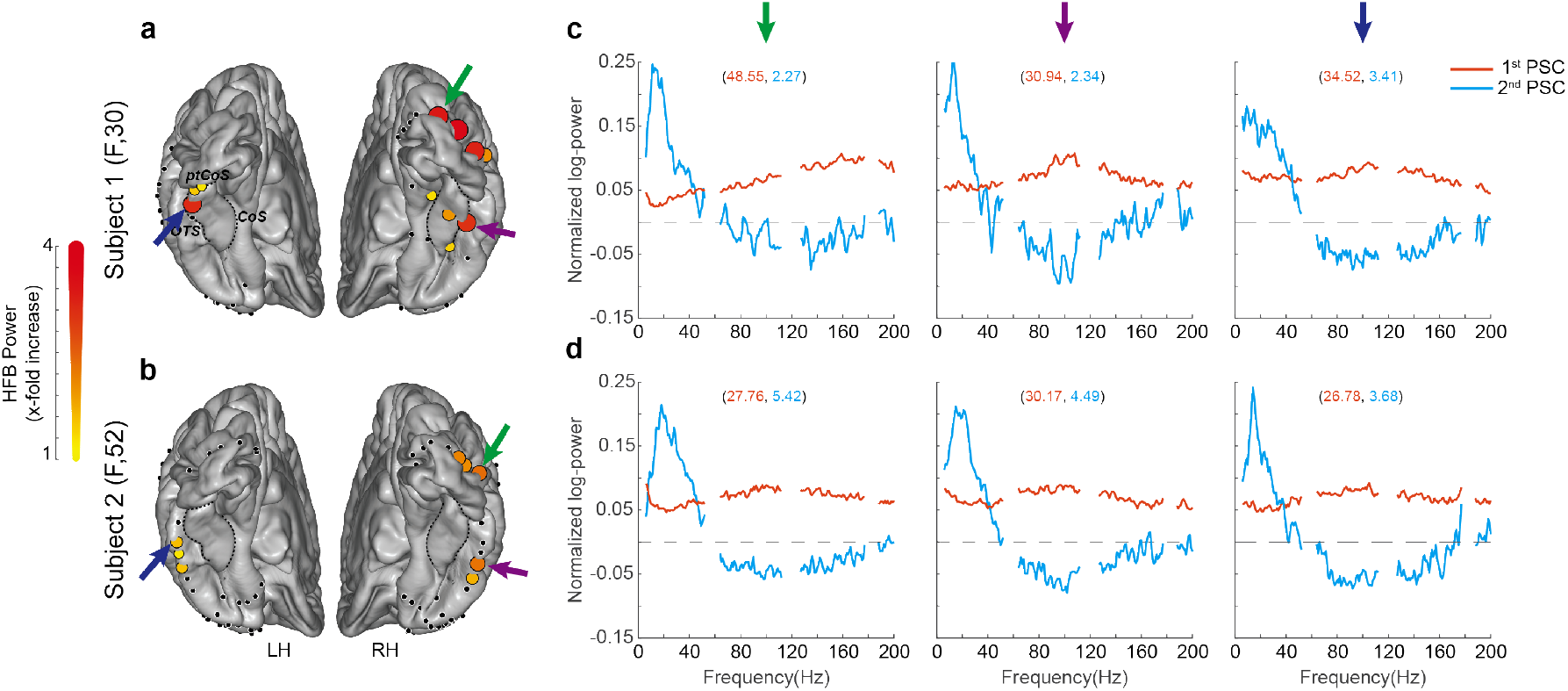
Power spectral components. (a,b) ECoG electrode locations in Subject 1 (top) and Subject 2 (bottom). Visual-responsive electrodes are shown in colored circles; color and size of the circle indicates the intensity of time-averaged HFB power increase (0-400 ms, face & word responses at 100% contrast). (c,d) First two PSCs for an EVC electrode (green arrow), a RH VTC electrode (purple arrow), and a LH VTC electrode (blue arrow) in each subject. The numbers in the parentheses denote the percent variance explained by each of the two PSCs. *COS* : collateral sulcus, *ptCOS* : posterior transverse COS, *OTS* : occipital-temporal sulcus

**Figure S3:**
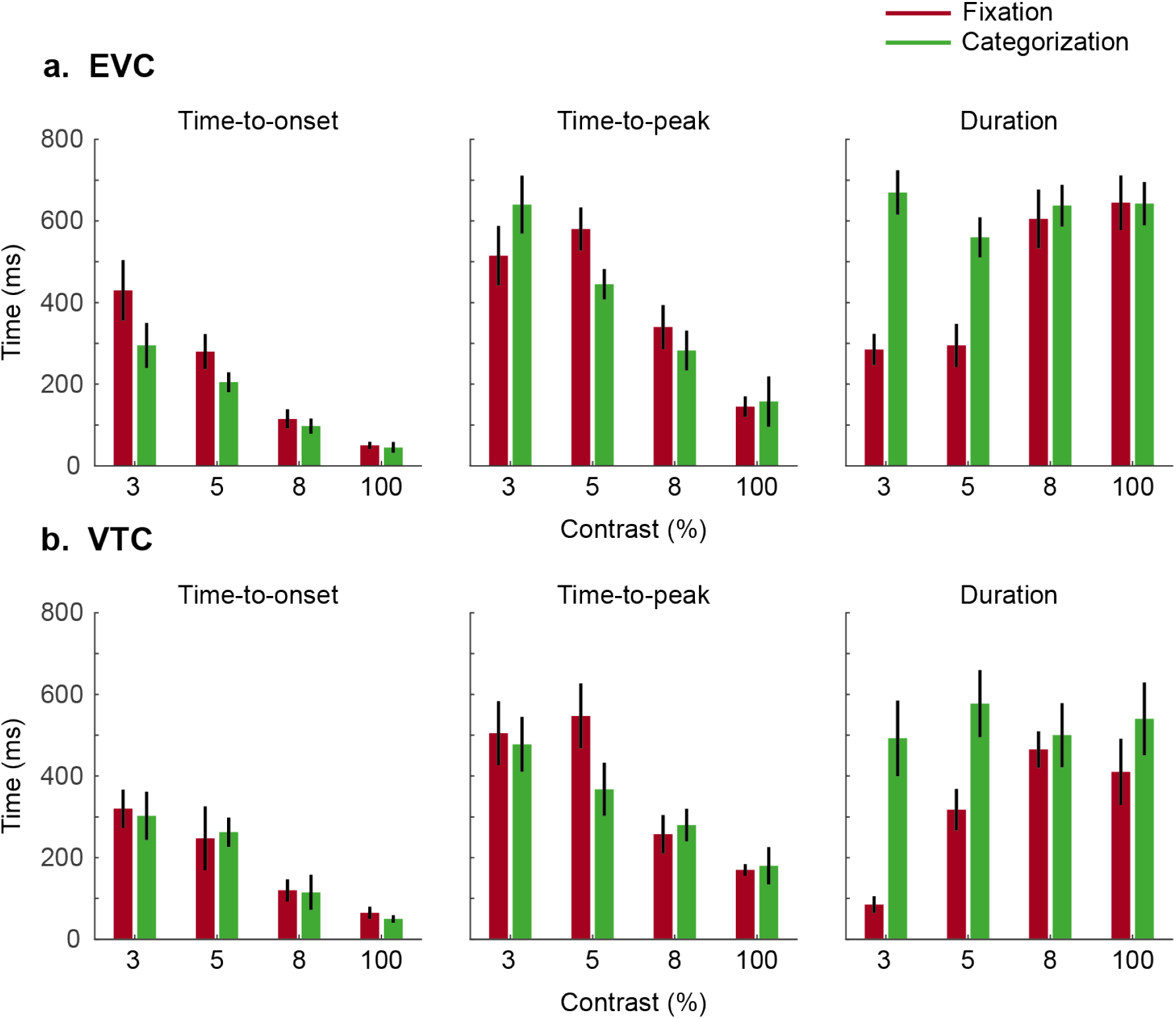
Temporal characteristics of HFB power increases. *Time-to-onset* (left), *time-to-peak* (center), and significant response *duration* (right) shown for (a) EVC and (b) VTC electrodes. Responses per condition were pooled for faces and words across EVC electrodes. For VTC electrodes, responses per condition were pooled only across the selective electrodes (i.e., faces from FFA electrodes and words from vWFA electrodes). Error bars indicate 68% confidence interval. Verticals bars indicate 68% confidence interval.

**Figure S4:**
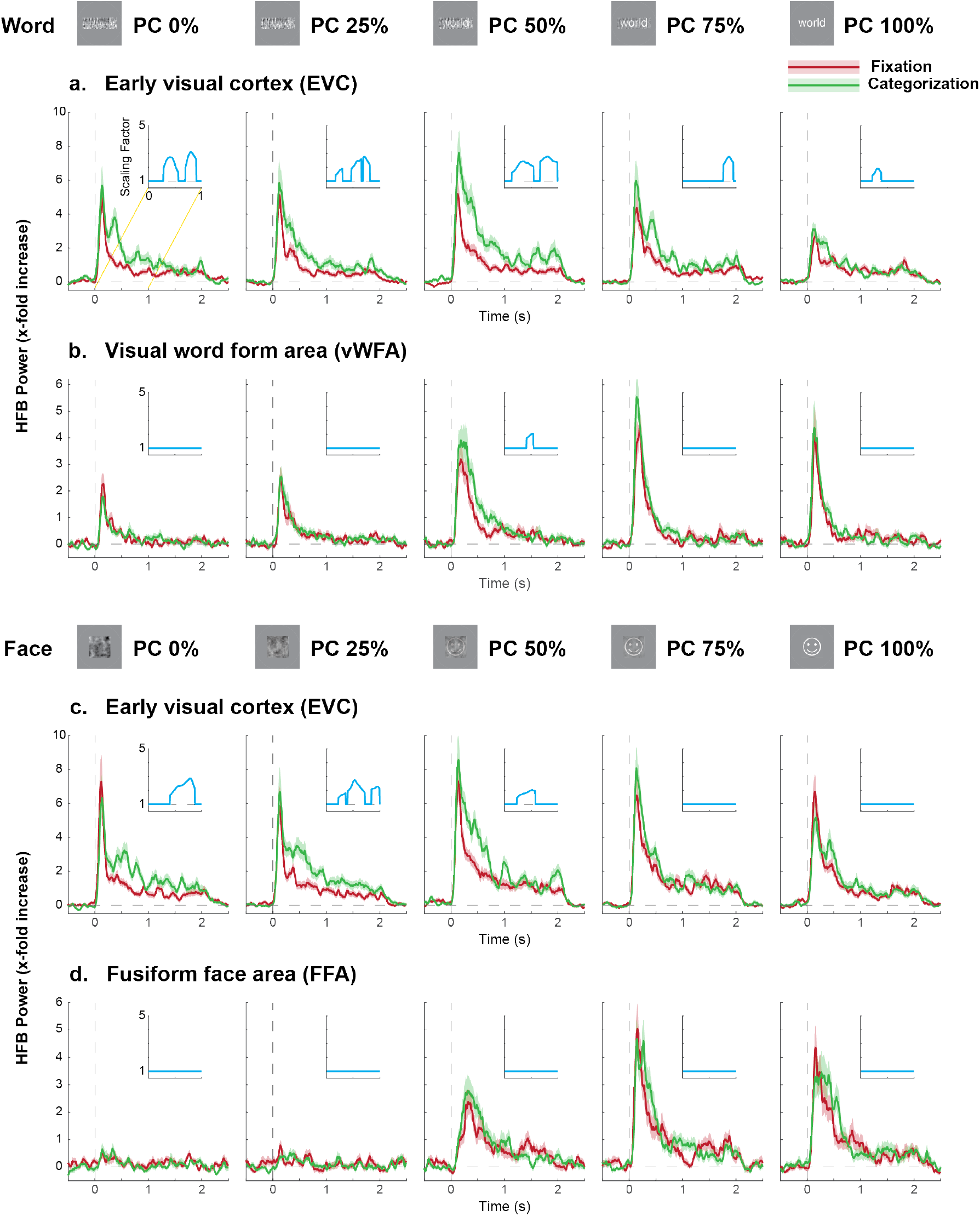
HFB power increases during phase coherence variation. Same as Figure 4, except for increasing stimulus phase coherence instead. ECoG HFB power traces during fixation (red) and categorization (green) with increasing stimulus phase coherence (word/face, shown on the top). The average HFB power (solid) was calculated by geometric averaging over electrodes and trials. The shaded region corresponds to the 68% confidence interval. The horizontal dashed lines indicate baseline and the vertical dashed lines indicate the stimulus onset. The inset in each plot corresponds to the scaling factor calculated as the ratio of HFB power of the categorization over the fixation task. Cyan traces deviated from 1 (grey dashed line) when scaling effects were significant (p<0.05; cluster-based permutation test). Dashed lines at 1 indicate no scaling.

**Figure S5:**
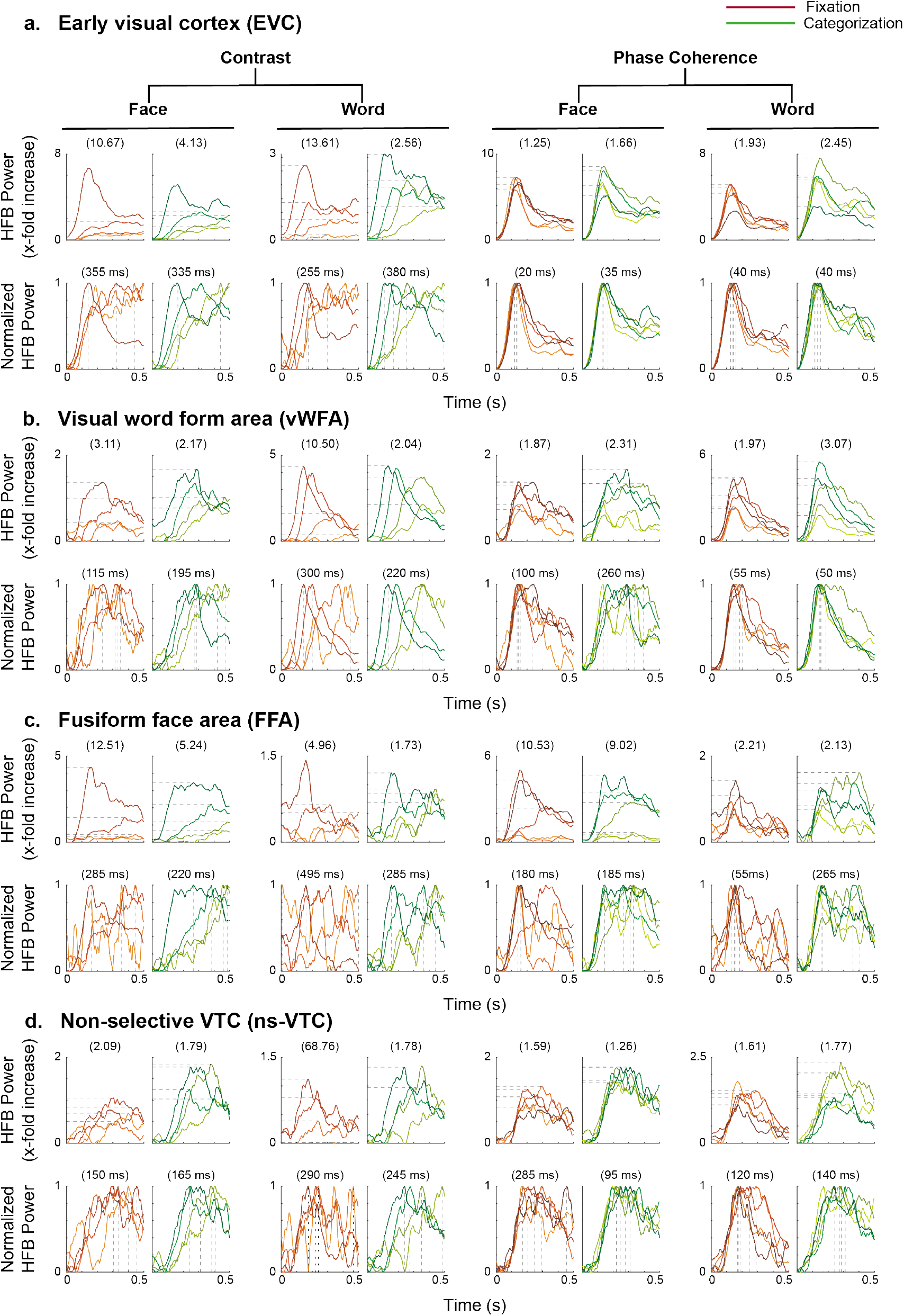
Summary plots of HFB power increases across stimulus manipulation. Top plots in each panel show average HFB power increases from 0 - 0.5 s for increasing contrast / phase coherence. The number in the parentheses indicate ratio of the peak HFB power (horizontal dashed lines) for the highest to the lowest contrast / phase coherence. Bottom plots in each panel show the normalized power. The number in the parentheses indicate the difference in the time to peak (vertical dashed lines) between the highest to the lowest contrast / phase coherence. Each colored line represents one stimulus condition with darker shades indicating higher contrast / phase coherence.

**Figure S6:**
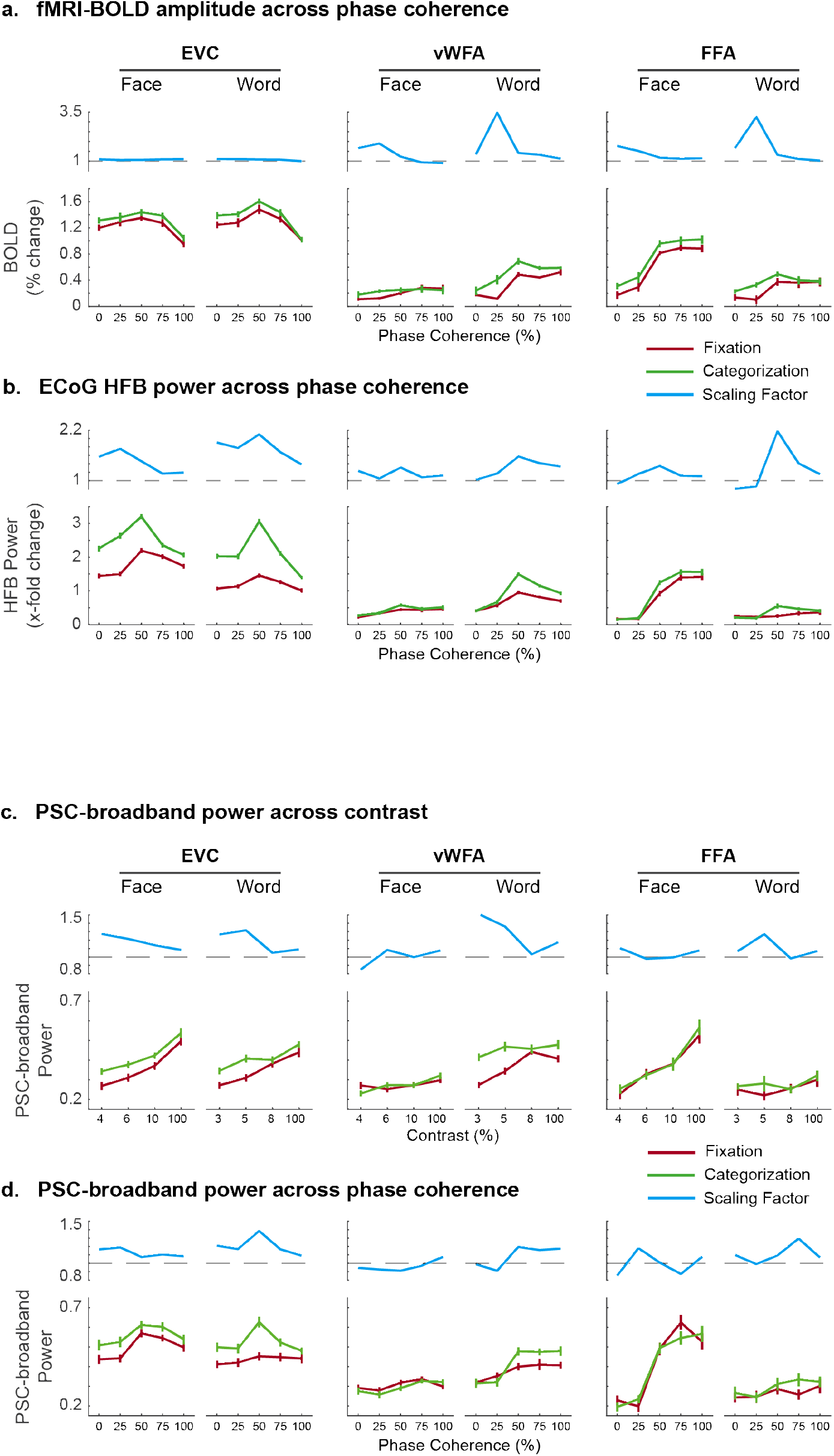
PSC-based HFB responses for contrast and phase coherence manipulations. (a,b) Time-averaged ECoG HFB responses match fMRI BOLD amplitudes for phase coherence manipulations. Same as Figure 5, except for variations across stimulus phase coherence instead. (c,d) HFB responses computed for varying stimulus (c) contrast and (d) phase coherence based on the first spectral PCA for EVC (left) vWFA (middle) and FFA (right) and time-averaged in 0-1 s time window. Error bars indicate 68% confidence interval. Cyan traces show scaling factors with horizontal dashed line at 1 indicating no scaling.

**Figure S7:**
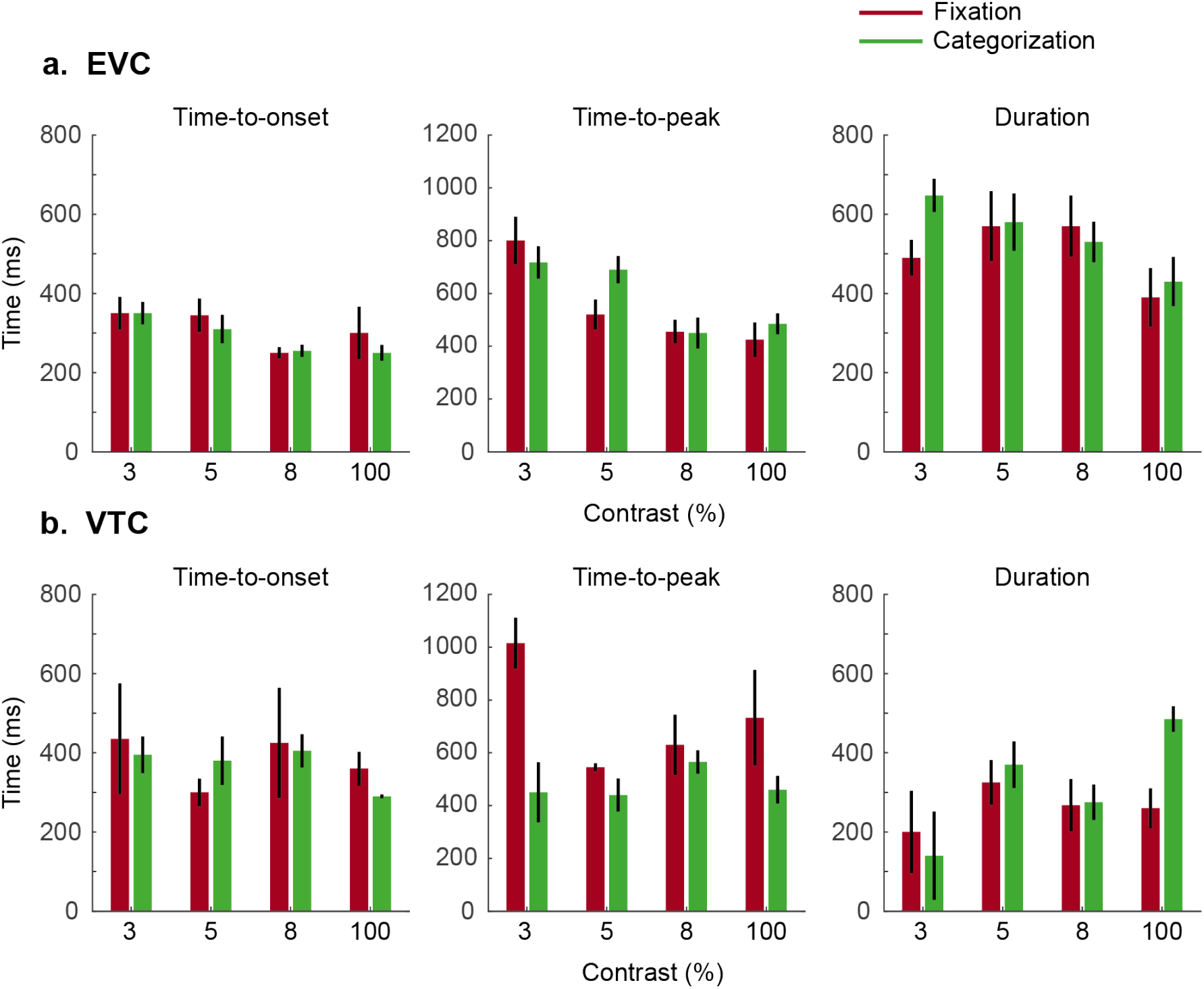
Temporal characteristics of LFB power decreases. *Time-to-onset* (left), *time-to-peak* decrease (center), and significant response *duration* (right) shown for (a) EVC and (b) VTC electrodes. Responses per condition were pooled for faces and words across EVC electrodes. For VTC electrodes, responses per condition were pooled only across the selective electrodes (i.e., faces from FFA electrodes and words from vWFA electrodes). Error bars indicate 68% confidence interval. Verticals bars indicate 68% confidence interval.

**Figure S8:**
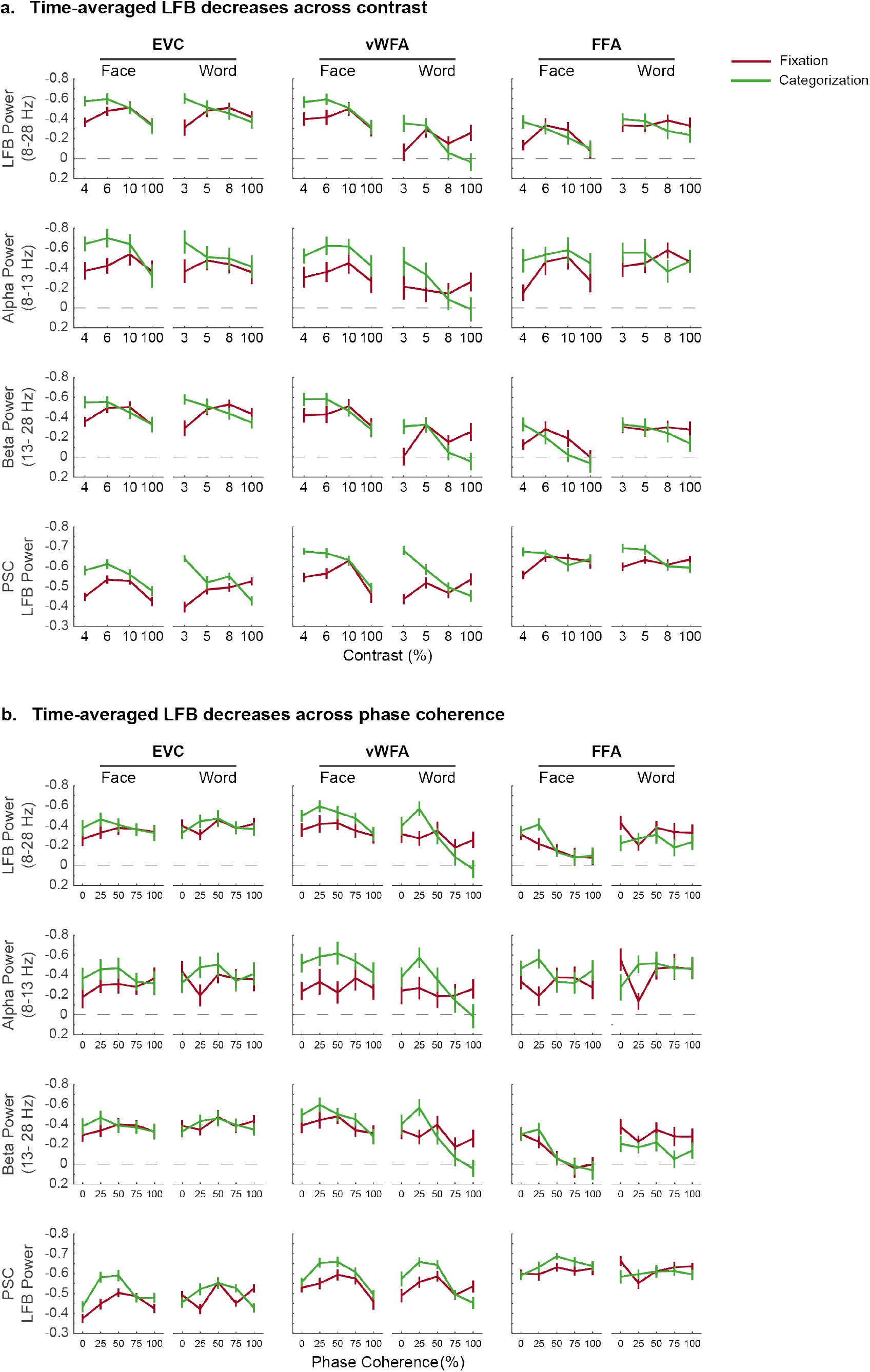
Time-averaged ECoG LFB decreases across different conditions. (a) ECoG responses time-averaged in 0.25-1.25 s time window using filtering in LFB (8-28 Hz; top), alpha (8-13 Hz, mid-top), and beta (13-18 Hz, mid-bottom) frequencies, and using second spectral PCA (bottom). Error bars indicate 68% confidence interval.

